# Handshake: Partner-Specific Protein-Protein Binding Site Prediction at Scale Using ProstT5 and Cross-Chain Attention

**DOI:** 10.64898/2026.06.04.730144

**Authors:** Nurit Haspel

## Abstract

Partner-specific protein-protein binding site prediction, identifying which residues of a protein form the interface when bound to a specific partner, remains a challenging task with significant implications for drug discovery and understanding of protein structure and function. Existing computational methods are limited by small training datasets, inconsistent redundancy filtering, and reliance on three-dimensional structural information at test time. Here we present a sequence-only, partner-specific protein-protein interface predictor called HandShake. It combines ProstT5, a protein language model pre-trained on structural data, with Low-Rank Adaptation (LoRA), a cross-chain attention mechanism and a contact supervision head. Our method can detect both binding interfaces and pairwise contact matrices. We trained our model on very large datasets of non-redundant protein-protein pairs derived from the PPInterface dataset, the most comprehensive structural protein-protein database to date, and evaluated it on systematically filtered benchmarks at four redundancy thresholds (30%–90% sequence identity). We demonstrate that sequence redundancy inflates reported AUROC by up to 0.079 and MCC by up to 0.145 on identical models, representing a substantial methodological confound in the field. Even at 30% redundancy threshold, our results (AUROC=0.811, MCC=0.367, F1=0.45) exceed the best published sequence-only result on this convention. Our method also achieves comparable performance to existing partner-specific methods that use explicit structural information. The comprehensive training and evaluation dataset, in addition to the systematic redundancy inflation, can help gain insight into protein-protein interactions and the abilities and limitations of current detection methods.

**Data availability:** The code and data can be found at http://github.com/nurith/Handshake.

**Author summary:** Understanding which residues of a protein make contact with a specific protein partner is fundamental for designing drugs and understanding cellular processes, but predicting these interfaces remains challenging. We developed a deep learning method that takes only the amino acid sequences of two proteins and predicts, for each residue, whether it lies on their binding interface. The method combines ProstT5, a protein language model trained to translate between sequence and structure, with a cross-chain attention mechanism that lets each protein’s residues attend to its partner. We train and evaluate on up to 32,503 non-redundant protein pairs across four sequence redundancy thresholds. To the best of our knowledge, this is by far the largest dataset trained and tested for this specific task. Our key methodological finding: sequence redundancy in evaluation benchmarks inflates reported metrics by up to 0.079 AUROC and 0.145 MCC on identical models when measured internally on PPInterface. Inference experiments on four independent benchmarks show that this internal inflation does not transfer cleanly to externally-curated datasets, where switching from 30%-trained to 90%-trained models changes performance only modestly. At 30% filtering, our method achieves AUROC values from 0.784 to 0.828 across five datasets, confirming genuine generalization. Through systematic negative controls and comparisons with the ESM-2 family of language models at matched parameter counts, we show that explicit structural pre-training, not just model scale, is what enables sequence-only binding site prediction to work.

## Introduction

Protein-protein interactions (PPIs) underlie the vast majority of cellular processes, and the specific residues mediating these interactions, the binding interface — determine the selectivity and affinity of the interaction. Predicting binding sites computationally is, therefore, of central importance for structural biology, drug design, and understanding disease mechanisms.

### Related Work

#### Protein Language Models for Interaction Prediction

ESM-2 [1] is a pure sequence-based transformer trained on ≈ 250 million protein sequences via masked language modeling, producing rich residue-level representations. ProstT5 [2] extends ProtT5-XL [3] by fine-tuning on structural data via Foldseek’s 3Di tokens, enabling translation between amino acid sequences and structural alphabets and endowing the model with explicit structural awareness without requiring structures at inference time. Several recent methods use ESM-2 to predict whether two proteins interact or not, training the methods on large PPI datasets. [4–6].

#### Protein-Protein Binding Site Prediction

Binding site prediction methods aim at predicting the specific region on the protein surface where proteins interact, when the protein structures are known. Two related but distinct formulations of this problem exist. Partner-independent prediction [7–11] asks which residues of an isolated protein are likely to interact with any other protein, while partner-specific prediction asks which residues form the interface between a specific protein pair. The latter is a harder and more informative task, as it requires the model to reason about both proteins simultaneously and predict complementary interfaces. Several partner-specific methods have been proposed, including BIPSPI [12], BIPSPI+ [13], PInet [14] and Pair-EGRET [15]. However, these methods share common limitations: they are trained and evaluated on small datasets (typically 230-3,972 complexes), they often require three-dimensional structural information at test time, and they employ inconsistent redundancy filtering strategies that may inflate reported performance. A separate line of work has focused on intrinsically disordered regions (IDRs). Disobind [16] uses ProtT5 embeddings for partner-specific intrinsically disordered residues (IDR) binding site and contact map prediction from sequence alone, but is restricted to IDR-containing complexes with fragments capped at 100 residues at training time (but not inference), representing a complementary rather than a competing approach. BIPSPI [12] introduced partner-specific prediction using XGBoost with sequence and structural features. BIPSPI+ [13] extended this work with larger, type-specific training datasets (up to ≈ 3, 972 complexes for heterodimers) and mixed redundancy filtering using SCOPe family pairs with 95% within-group identity and 30% identity for unclassified proteins, achieving sequence-only MCC of 0.279—0.311 depending on the dataset. Pair-EGRET [15] uses edge-aggregated graph attention networks with protein language model features and cross-chain attention on three benchmark datasets (DBD5 [17]: 230 complexes; Dockground [18, 19]: 396 complexes; MaSIF [20]: 3,147 complexes filtered for length and contact percentage). Crucially, Pair-EGRET requires three-dimensional structural coordinates at both training and test time, making it unsuitable for proteins lacking experimental structures. PInet [14] represents the proteins as pairs of point clouds encoding the structures of two partner proteins, capturing both geometric and physicochemical molecular surface complementarity. PAIRpred [21] is an earlier SVM-based method that uses pairwise kernels on sequence and structural features, evaluated on Docking Benchmark 4.0. It established much of the foundational methodology for partner-specific prediction.

#### Redundancy in PPI Benchmarks

Sequence redundancy in training and test sets is a known confounding factor in machine learning benchmarks for bioinformatics. When training and test proteins share high sequence identity, models can achieve inflated performance by memorizing homologous interfaces rather than learning general binding site geometry. Despite this, most published partner-specific methods do not systematically report the effect of redundancy threshold on performance, making cross-study comparisons unreliable.

#### Our Contribution

Here we address these limitations by presenting a partner-specific binding site predictor that: (1) operates from sequence alone at inference time; (2) is trained on very large datasets: 31,000 non-redundant protein pairs, up to nearly 32,000 protein pairs at 90% sequence redundancy. *To the best of our knowledge, it is the largest dataset ever used for this task*; (3) employs systematic redundancy filtering across four identity thresholds, allowing us to quantify redundancy inflation directly; and (4) uses ProstT5, a protein language model explicitly pre-trained on structural data via Foldseek’s 3Di structural alphabet, providing structural awareness without requiring structures at test time. At rigorous 30% identity filtering, our method achieves AUROC=0.811 and MCC=0.367; permissive 90% filtering inflates these to AUROC=0.890 and MCC=0.512. A three-tier hierarchy of negative controls confirms the model learns real sequence-to-binding-site signal, and size-matched comparisons with ESM-2 demonstrate that ProstT5’s structural pre-training accounts for the bulk of its advantage on this task across all distance conventions tested.

## Materials and methods

### Dataset

We derived our dataset from the PPInterface database [22]. The entire dataset contains over 800,000 entries. After removing all the non-existent or non-standard entries, all pairs with *<* 1% or *>* 80% interface residues, and limiting the chain sizes to 50–300, we retained 233,538 raw pairs. We then generated four non-redundant subsets using CD-HIT [23] at 90%, 70%, and 50% sequence identity thresholds, and MMseqs2 [24] at 30% (CD-HIT does not reliably handle thresholds below approximately 40%). Non-redundancy filtering was performed on concatenated pair sequences (chain A + chain B), following the approach of [12], retaining one representative per cluster.

Binding site labels were assigned using a 6Å heavy-atom distance threshold, matching the convention used by several published methods including BIPSPI+ [13] and Pair-EGRET [15]: residue *i* in chain A is labeled as binding (1) if any non-hydrogen atom of residue *i* is within 6Å of any non-hydrogen atom of any residue in chain B, and 0 otherwise. We recalculated the interfacing residues, since it is a slightly different convention to the one used by PPInterface [22]. The resulting class imbalance is approximately 13% binding residues, between the otherwise used distance conventions of 6Å *Cα*–*Cα* (with ≈ 5% binding residues) and 8Å *Cα*–*Cα* (with ≈ 17% binding residues). The difference in performance between the three distance conventions is reported in Supplementary Tables 3–7. After filtering, the four datasets contain 14,005 pairs (30% sequence redundancy using MMseqs2), 18,146 pairs (50% CD-HIT), and 32,503 pairs (90% CD-HIT). The pair count grows non-linearly with redundancy threshold: the 30%–50% step adds only 4,141 pairs (30%), while the 50%–90% step adds 14,357 pairs (79%), reflecting the increasing fraction of homologous pairs at higher identity thresholds. For all experiments, pairs were split 70/15/15 into training/validation/test sets using a fixed random seed (42), ensuring identical splits across all redundancy levels for fair comparison. Pairs are de-duplicated under chain reordering (i.e., `1ABC_A_1ABC_B` and `1ABC_B_1ABC_A` are treated as the same pair) prior to splitting, eliminating any train/test overlap from this source.

### Model Architecture

The model pipeline is depicted in Figure 1. Our model combines three components: a frozen-then-LoRA-tuned ProstT5 encoder, a cross-chain attention module, and two prediction heads. We denote the two input chains as *A* and *B*, with lengths *L*_*A*_ and *L*_*B*_, respectively, and use *d* = 1024 for the hidden dimension of ProstT5.

**Fig 1.**
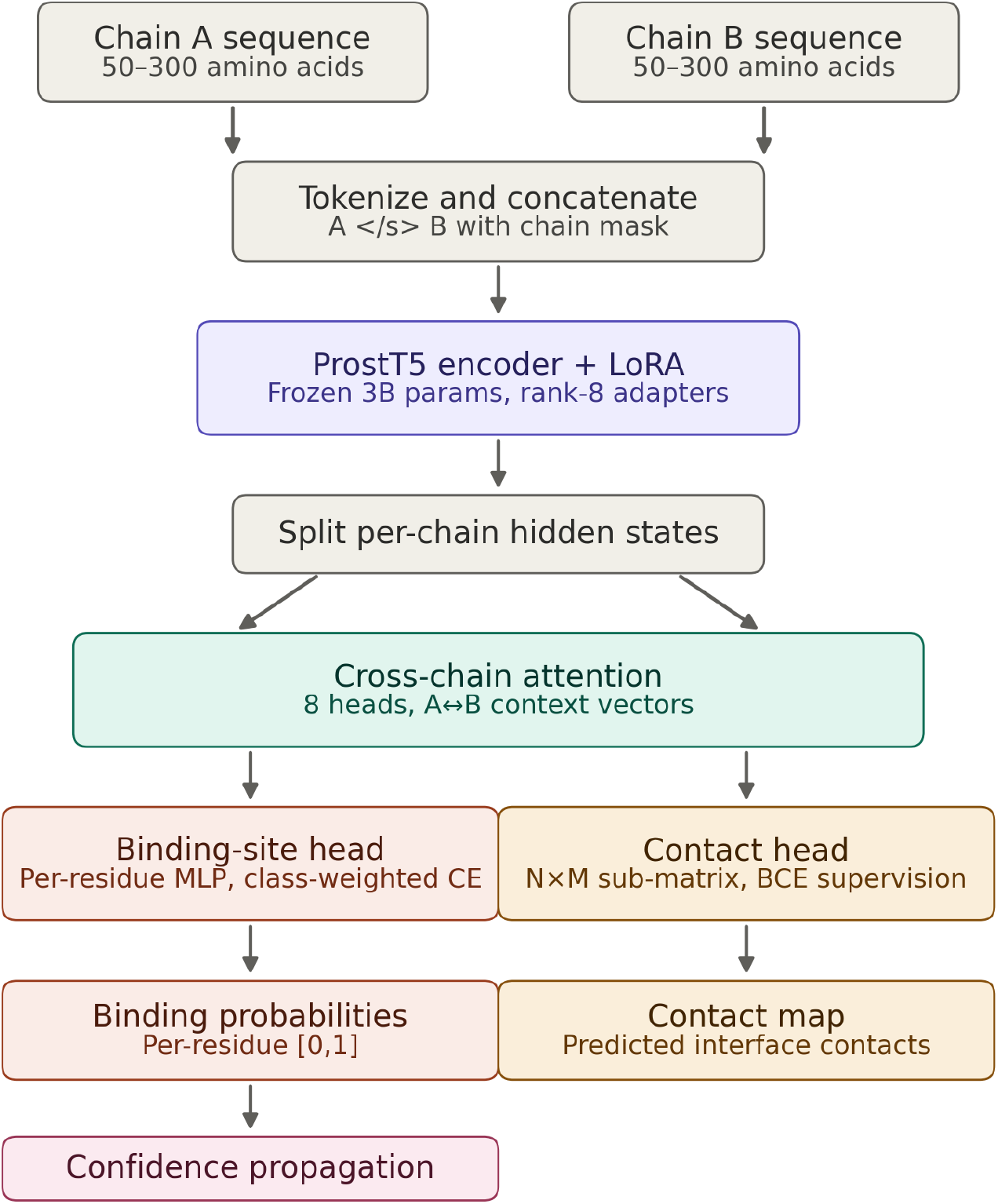
Overview of the pipeline. The input is two protein sequences with per-residue labels. The chains are tokenized and concatenated with a separator indicating the split point. The input is encoded with the ProstT5 encoder using a rank-8 Low-Rank Adaptation. A multi-head cross-chain attention mechanism computes attention between the two chains. The binding-site head and the contact head compute per-residue binding probabilities and the interface contact map, respectively. An optional confidence-propagation step can be applied to enhance F1 or MCC values.

#### ProstT5 Encoder with LoRA

The initial architecture was based on the public ProtTrans github [3] and was heavily modified to accommodate the task at hand. ProstT5 (based on ProtT5-XL, ≈ 3*B* parameters) is used as the sequence encoder. Both chains are space-tokenized using the ProstT5 tokenizer and concatenated with a ⟨/s⟩ separator token, yielding a single input sequence of length *L*_*A*_ + *L*_*B*_ + 1. After the encoder forward pass the hidden states 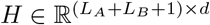 are split back into per-chain representations 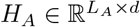 and 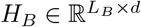. We apply the low-rank adapter formulation used in the public ProtTrans fine-tuning code [3], which combines an additive low-rank update (as in LoRA [25]) with a multiplicative rank-1 scaling. We use rank *r* = 8 applied to the query and value projection matrices of every encoder attention layer, freezing all other encoder parameters. For each adapted weight matrix 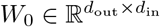, the effective weight is:

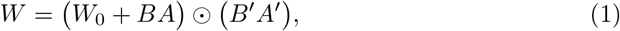

where 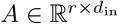 with entries initialized as 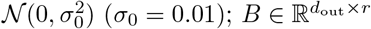 initialized to zero so the additive adapter starts as a no-op; and 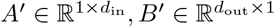 initialized to all-ones plus small Gaussian perturbations, so the multiplicative scaling starts as approximately the identity. The symbol ⊙ denotes elementwise multiplication after broadcasting the rank-1 multiplicative term to the shape of *W*_0_ + *BA*. Only *A, B, A*′, *B*′ are trained; *W*_0_ remains frozen. The ablation labeled “LoRA *r* = 0” disables both mechanisms (*BA* = 0 and *B*′*A*′ = 1), giving a fully frozen encoder.

#### Cross-Chain Attention

A multi-head cross-chain attention module (*h* = 8 heads, hidden size *d* = 1024) takes the per-chain ProstT5 representations and computes cross-attention weights between chain A residues (queries) and chain B residues (keys/values), and vice versa. We use the standard scaled dot-product attention formulation [26]. For each direction, with per-head dimension *d*_*h*_ = *d/h*:

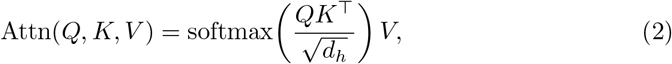

where *Q* = *H*_*A*_*W*_*Q*_, *K* = *H*_*B*_*W*_*K*_, *V* = *H*_*B*_*W*_*V*_ for the A → B direction, and symmetrically for B → A. The resulting context vectors 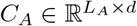 and 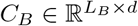 summarize each residue’s expected partner context.

We additionally retain the raw (pre-projection) attention weight matrices 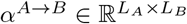 and 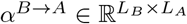 for use in the contact head and attention-supervision loss described below. This module is the primary mechanism by which partner context is incorporated into per-residue predictions.

#### Binding Site Head

Each residue receives a final representation concatenating its ProstT5 embedding and its cross-attention context vector. A two-layer multi-layer perceptron *g*_*θ*_ (hidden sizes 64 and 32, with ReLU activations) produces per-residue binding-site logits:

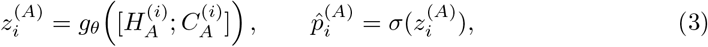

and symmetrically for chain B. Here [;] denotes concatenation and *σ* the sigmoid (after softmax over the two-class logit).

#### Contact Head

A secondary contact head produces an *L*_*A*_ × *L*_*B*_ residue-residue contact matrix. We first construct a pairwise feature tensor by summing the two attention directions and adding a small MLP applied to the outer combination of cross-attention weights:

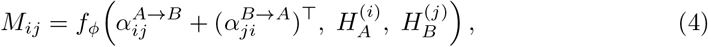

where *f*_*ϕ*_ is a small MLP. Predicted contact probabilities *ĉ*_*ij*_ = *σ*(*M*_*ij*_) are computed only within the sub-matrix of residues whose predicted binding probabilities exceed a threshold *τ*_bind_ = 0.3, i.e. over 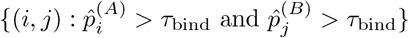.

#### Loss Function

The total training loss combines three terms: a per-residue binding-site loss, a contact-map loss, and an attention-supervision loss:

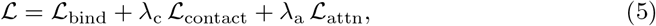

with *λ*_c_ = 1.0 and *λ*_a_ = 0.5. The binding loss is a class-weighted cross-entropy with label smoothing *α*_LS_ = 0.05 and class weights *w*_0_ = 1 (non-binding), *w*_1_ = 3 (binding):

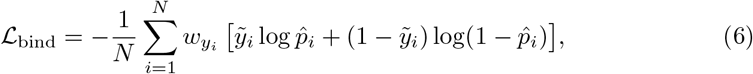

where 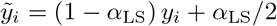 is the smoothed label and the sum runs over all *N* valid residues in the batch (both chains, excluding padding).

The contact loss is a binary cross-entropy over the predicted-binding sub-matrix, with a per-pair positive weight 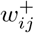 capped at *w*_max_ = 10 to limit gradient blow-up on extremely sparse positives:

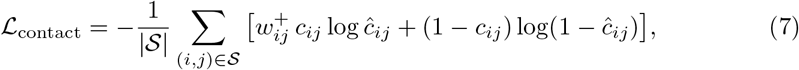

where 𝒮 is the predicted-binding sub-matrix and *c*_*ij*_ is the ground-truth contact label.

The attention-supervision loss guides the raw cross-attention weights toward known contacts using Kullback–Leibler divergence between the row-normalized attention distribution and the row-normalized true-contact distribution:

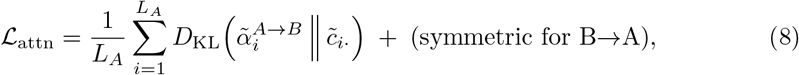

where 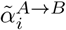 is the softmax-normalized row of attention weights and 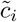 is the row-normalized ground-truth contact vector. Rows with no ground-truth contacts are excluded from the sum.

#### Training

Models were trained for 12 epochs using the AdamW optimizer [27] with hyperparameters *β*_1_ = 0.9, *β*_2_ = 0.999, *ϵ* = 10^−8^, and weight decay *λ*_WD_ = 0.01. A cosine learning-rate schedule was used with a peak learning rate of *η*_max_ = 3 × 10^−4^ and 100 linear warmup steps, decaying smoothly to zero by the final step:

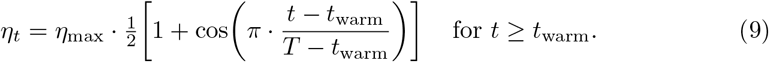

The per-device batch size was 4 with gradient accumulation over 8 steps, giving an effective batch size of 32 pairs. Training used mixed-precision (bfloat16) arithmetic on a single NVIDIA A100-SXM4 40GB GPU. Gradients were clipped to a global L2 norm of 1.0. The best checkpoint was selected based on validation MCC.

#### Threshold Tuning and Confidence Propagation

The default prediction threshold of *τ* = 0.5 was tuned by sweeping the validation set for either MCC (default) or F1 maximization:

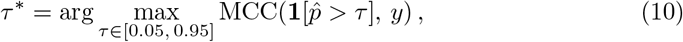

with *τ* swept in steps of 0.01.

Post-hoc confidence propagation was applied to the binary predictions to enforce spatial coherence of binding patches. Residues with mid-range probabilities *p*_*i*_ ∈ (*τ*_lo_, *τ*_hi_) are promoted to binding if they have at least *k*_min_ predicted-binding neighbors within a 1D sequence window of half-width *w*; isolated core binding residues with fewer than *k*_core_ predicted-binding neighbors can be demoted. Formally, let 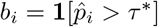 and *n*_*i*_ = ∑_|*j*−*i*|≤*w, j* ≠ *i*_ *b*_*j*_. The propagated prediction is:

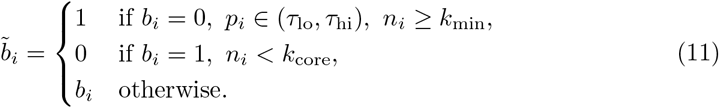

Optimal parameters (*τ*_lo_, *τ*_hi_, *w, k*_min_, *k*_core_) were selected by grid search over thousands of combinations on the validation set for MCC and F1.

### Evaluation Metrics

We report several complementary metrics for binary classification. Let TP, FP, TN, FN denote the standard confusion-matrix counts on the test set, computed at the per-residue level across all residues of both chains in all test pairs. Precision, recall, and the *F*_1_ score are:

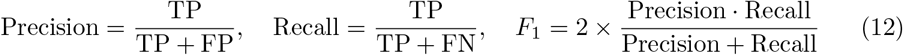

Matthews’ correlation coefficient [28] (MCC) is a balanced measure that takes all four confusion-matrix quantities into account:

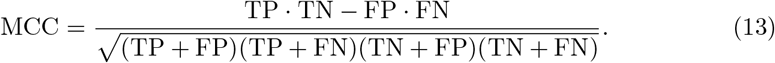

The MCC ranges from −1 (perfectly wrong) to 0 (random) to +1 (perfect) and unlike *F*_1_ does not collapse to artificially high values when one class dominates the predictions.

AUROC and AUPRC are computed from the predicted probabilities 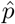 without thresholding. AUROC is the area under the receiver-operating-characteristic curve and equals the probability that a randomly drawn positive example is ranked above a randomly drawn negative example. AUPRC is the area under the precision–recall curve; its random baseline equals the positive-class prevalence *π* = 𝔼[*y*], so we additionally report AUPRC lift defined as AUPRC*/π* for cross-dataset comparison. AUROC and AUPRC are our primary threshold-independent metrics, alongside MCC, *F* 1, precision, and recall at tuned thresholds. AUROC is preferred for cross-method comparison since it is reported by most published partner-specific methods (BIPSPI, Pair-EGRET, MaSIF) and is less sensitive to the extreme class imbalance of the task (≈ 13% binding). For contact prediction, we report AUPRC computed within the predicted-binding sub-matrix at *τ*_bind_ = 0.3, comparing against a random baseline equal to the contact density within the sub-matrix.

## Results

### Baseline and Control

#### Negative Control

To verify that the model learns genuine sequence-to-binding-site signal rather than dataset artifacts, we trained two negative-control variants (1) **Globally random labels**: labels are drawn independently from a Bernoulli distribution at the empirical binding rate.(2) **Within-chain shuffled labels**: binding labels are randomly permuted within each chain, preserving per-chain binding rate but breaking the within-chain sequence-to-label relationship. The results are shown in Table S1 and Figure S1. With globally random labels (independent Bernoulli sampling at the empirical binding rate), our model achieved AUROC=0.500 and MCC=0.001, which is essentially a random level. All but 11 of 681,827 residue pairs were predicted as non-binding, confirming no information leakage in the data pipeline or model architecture. Changing labels within the chain (preserving the per-chain binding rate but breaking the position-specific signal) yield AUROC=0.653 and MCC=0.161, meaningfully above random, reflecting the signal of the per-chain composition that is real biology (the binding percentage varied among pairs. For example, chains with more aromatic residues tend to contain more interface residues). As shown below, The full model with real labels achieves AUROC=0.811 on the same dataset, an additional +0.158 AUROC above the shuffled baseline. This three-tier hierarchy decomposes the model’s performance into three distinct sources of signal.

#### Architectural Ablation

We performed ablation experiments to assess the contribution of cross-attention and contact supervision, as well as the role of LoRA fine-tuning (Table S2). Results are reported with both LoRA rank 8 (fine-tuned encoder) and LoRA rank 0 (truly frozen encoder). The two mechanisms, cross-chain attention and LoRA fine-tuning, are remarkably substitutable. Removing cross-attention costs +0.065 MCC when the encoder is frozen (LoRA=0) but only +0.025 MCC when the encoder is fine-tuned (LoRA=8). Symmetrically, removing LoRA costs +0.048 MCC without cross-attention but only +0.007 MCC with it. Most strikingly, the no-cross-attention LoRA=8 configuration (MCC=0.343, AUROC=0.793) is only slightly inferior to the full-model LoRA=0 configuration (MCC=0.360, AUROC=0.802): two architecturally distinct mechanisms, LoRA adapting the encoder representations vs cross-chain attention combining frozen representations, arrive at a similar final performance. This result has a practical consequence: the frozen ProstT5 model with cross-attention (MCC=0.360, F1=0.439, AUROC=0.802, AUPRC=0.451) achieves nearly the same performance as the fully fine-tuned model (MCC=0.367, F1=0.444, AUROC=0.811, AUPRC=0.462) at substantially lower computational cost, an attractive option for resource-constrained deployment.

#### Comparison With ESM-2

To assess the contribution of ProstT5’s structural pre-training, we trained ESM-2 650M and ESM-2 3B with the identical cross-chain attention architecture on the dataset. ESM-2 3B provides a size-matched comparison with ProstT5 (both 3*B* parameters), so any performance difference reflects the encoder pre-training objective rather than model capacity. With frozen encoders, ESM-2 3B achieves MCC=0.277 and AUROC=0.754, while our full ProstT5 model (with LoRA fine-tuning) achieves MCC=0.367 and AUROC=0.811, a size-matched gap of +0.090 MCC, +0.057 AUROC, and +0.101 AUPRC favoring ProstT5. Scaling ESM-2 from 650M to 3B improves MCC by only +0.033, indicating that ESM-2 saturates with model size on this task and the gap is not closed by additional capacity. LoRA fine-tuning has opposite effects on the two encoders. For ProstT5, LoRA improves performance, albeit slightly in the presence of cross-attention. For ESM-2, LoRA fine-tuning consistently destabilized training across all distance conventions tested (Table S8), producing complete or near-complete collapse to the majority class regardless of class weight, rank, or contact-supervision settings. This asymmetry is striking: the same fine-tuning approach that gently improves a structurally pre-trained encoder destroys a sequence-only encoder, suggesting that ProstT5’s 3Di pre-training creates a loss landscape amenable to task-specific adaptation while ESM-2’s purely sequence-based representations are disrupted by gradient updates.

#### Redundancy Inflation

We first quantify how sequence redundancy in the training and test sets affects reported performance. Using identical model architecture and hyperparameters across four redundancy thresholds, we observe a monotonic increase in all performance metrics as the filtering threshold is relaxed (Table 1 and Figure 2). MCC increases from 0.367 at 30% identity to 0.512 at 90%, a difference of +0.145 MCC entirely attributable to evaluation rigor rather than model quality. AUROC increases from 0.811 to 0.890 (+0.079), AUPRC from 0.462 to 0.624 (+0.144), and F1 from 0.452 to 0.568 (+0.116). Beyond the absolute inflation, the curve shape is informative. The 30%→50% step adds 4,141 pairs (≈ 30% increase) for +0.075 MCC – a high per-pair efficiency of 0.018 MCC per 1,000 pairs, consistent with new pairs contributing genuinely diverse signal. The 50% →90% step then adds 14,345 pairs (nearly 80% increase) for +0.07 MCC – 0.0049 MCC per 1,000 pairs, showing that the new pairs are more homologous and therefore less informative biologically. This dissociation between dataset diversity and apparent performance gain isolates homolog memorization as the dominant mechanism behind the 50%-to-90% performance jump.

**Table 1.** Performance across sequence redundancy thresholds following MCC-threshold tuning + confidence propagation).

| Dataset | Pairs | MCC | F1 | AUROC | AUPRC | Precision | Recall |
| --- | --- | --- | --- | --- | --- | --- | --- |
| 30% (MMseqs) | 14,005 | 0.367 | 0.452 | 0.811 | 0.462 | 0.46 | 0.43 |
| 50% (CD-HIT) | 18,146 | 0.442 | 0.517 | 0.854 | 0.536 | 0.40 | 0.51 |
| 90% (CD-HIT) | 32,500 | 0.512 | 0.568 | 0.890 | 0.624 | 0.57 | 0.57 |

**Fig 2.**
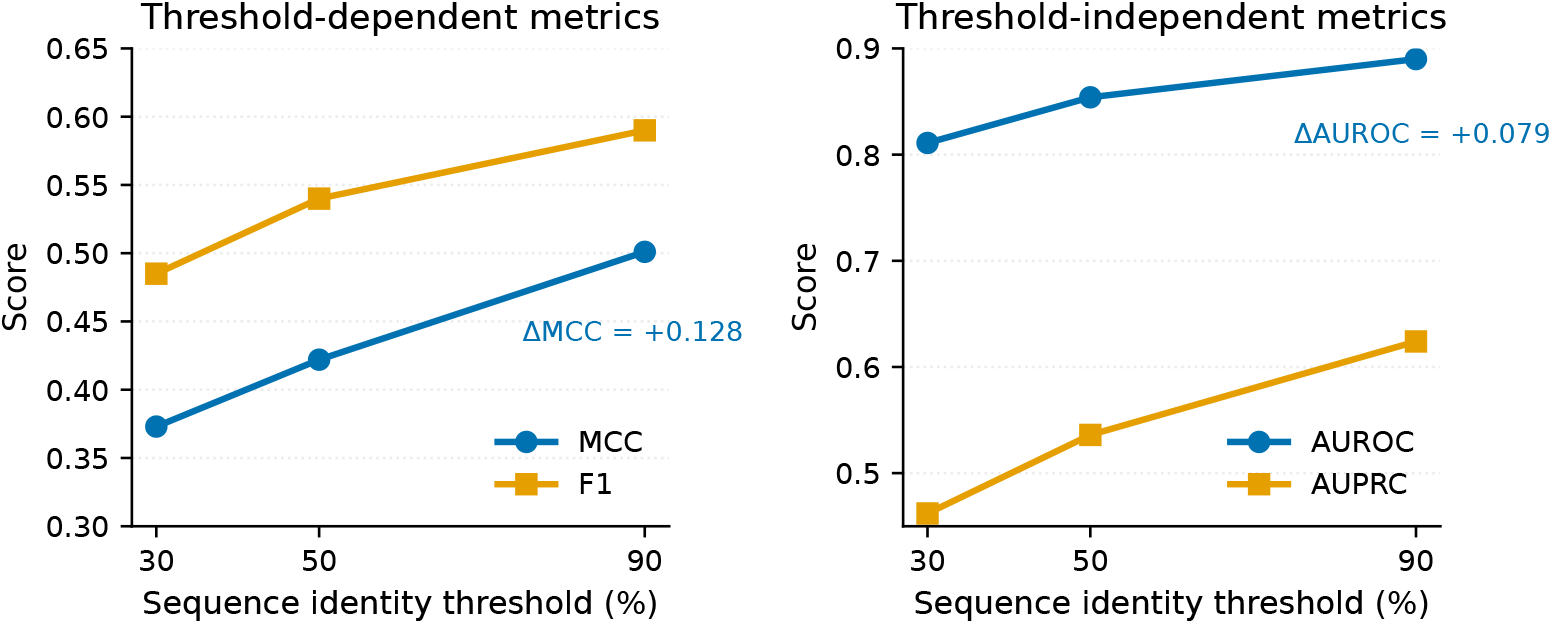
Evaluation metrics as a function of PPInterface dataset sequence redundancy (30%-90%), as shown in Table 1 Left: Threshold dependent values. Right: Threshold-independent values. In both cases, there is a sharp increase as sequence redundancy grows.

### Comparison with Published Methods

Direct numerical comparison with published methods is complicated by differences in datasets, redundancy filtering strategies, distance thresholds, and evaluation metrics. With these caveats, our 30% MMseqs2 result (MCC=0.373, AUROC=0.789) is competitive with or exceeds sequence-only partner specific published methods on the same convention. BIPSPI+ [13] reports sequence-only MCC of 0.279–0.311 on heterodimer datasets of 2,401–3,972 complexes with mixed redundancy filtering (30%–95% depending on SCOPe family availability); our MCC=0.373 exceeds this on a substantially larger (≈ 4×) and more systematically non-redundant benchmark. Among partner-independent methods evaluated on the standard Test 60 benchmark, the best sequence-only method (EnsemPPIS [10]) reports MCC=0.266, and the best structure-based methods such as EquiPPIS [8] and MIPPIS [9] report MCC≈0.37. While these numbers are not directly comparable to ours (different datasets, partner-independent vs. partner-specific task), they provide context: our sequence-only partner-specific method achieves metrics in the range of the best published structure-based partner-independent methods, while solving a harder task variant without requiring 3D coordinates at test time.

#### Testing on Benchmark Datasets

To assess generalization beyond PPInterface and compare to other partner-specific methods trained and tested on much smaller datasets, we measured the performance of our model on several available datasets: MaSIF, DBD5, Dockground Benchmark 4 and EpIPred. External datasets often contain multimeric complexes. Since our architecture accepts pairs of proteins, trimeric entries such as AB C were split into the constituent dimeric interfaces A C and B C (but not A B), while entries with ambiguous multi-chain interfaces (e.g., AB CD) were excluded. Interface residues were re-labeled using our 6Å heavy-atom convention for consistency across all datasets. This decomposition and re-labeling means that our evaluation sets are not identical to those used by published methods. For example, our DBD5 evaluation set contains 335 pairs derived from the original 230 complexes, after excluding multi-chain entries that could not be unambiguously decomposed into binary interfaces. Direct numerical comparison with methods evaluated by other methods should be considered with this caveat.

We trained and tested our architecture from scratch on the MaSIF dataset, as well as a non-redundant subset 3,979 pairs, after filtering for 30% sequence redundancy. The other datasets are too small to give meaningful results on our architecture. We therefore performed inference-only experiments using the PPInterface-trained model at 30% and 90% filtering thresholds (Table 2).

**Table 2.** Cross-dataset experiments. MaSIF, DBD5, EpIPred, Dockground, with comparison to other methods when available.

| Dataset | Model | AUROC | AUPRC | MCC | Precision | Recall |
| --- | --- | --- | --- | --- | --- | --- |
| Dockground | Handshake (inf. 30%)<sup>†</sup> | 0.802 | 0.411 | 0.342 | 0.38 | 0.49 |
|  | Handshake (inf. 90%)<sup>‡</sup> | 0.816 | 0.443 | 0.371 | 0.43 | 0.47 |
| Epipred | Handshake (inf. 30%) | 0.784 | 0.393 | 0.323 | 0.37 | 0.47 |
|  | Handshake (inf. 90%) | 0.799 | 0.422 | 0.351 | 0.43 | 0.43 |
|  | PInet | 0.751 | 0.330 | – | 0.19 | 0.90 |
| MaSIF (30%) | Handshake | 0.828 | 0.585 | 0.438 | 0.54 | 0.57 |
| MaSIF (full) | Handshake | 0.834 | 0.575 | 0.435 | 0.54 | 0.54 |
|  | PInet [14] | 0.88 | 0.45 | – | 0.31 | 0.63 |
|  | Pair-EGRET [15] | 0.958 | 0.594 | – | – | – |
| DBD5 | Handshake (inf. 30%) | 0.809 | 0.436 | 0.359 | 0.40 | 0.49 |
|  | Handshake (inf. 90%) | 0.842 | 0.511 | 0.416 | 0.55 | 0.41 |
|  | BIPSPI (seq only) | 0.753 | 0.305 | 0.279 | 0.30 | 0.48 |
|  | BIPSPI | 0.822 | 0.410 | 0.385 | 0.39 | 0.56 |
|  | BIPSPI+ | 0.848 | 0.465 | – | 0.44 | 0.57 |
|  | PInet | 0.837 | 0.667 | 0.459 | 0.51 | 0.75 |
|  | Pair-Egret | 0.924 | 0.275 | – | 0.26 | 0.75 |

<sup>†</sup> Inference only with the weights from the PPInt dataset filtered at 30%. It is the same weights as the model shown in Table 1
<sup>‡</sup> Inference only with the weights from the PPInt dataset filtered at 90%. It is the same weights as the model shown in Table 1

The MaSIF dataset [20] contained 5,523 pairs. On the full dataset, the model achieves MCC=0.435 and AUROC=0.834. After applying 30% identity filtering (3,979 pairs), performance remains essentially unchanged, at MCC=0.438 and AUROC=0.828. Strikingly, performance at 30% filtering on MaSIF exceeds the performance on PPInterface, confirming that our method generalizes across independently curated PPI databases when redundancy is properly controlled and that the architecture is not overfit to PPInterface-specific characteristics. Our method achieves lower AUROC but otherwise comparable performance to PInet and Pair-EGRET, despite the latter two methods using direct structural information. We did not use the MaSIF’s predefined train/test split as appearing on their github. It yielded substantially lower performance (AUROC=0.747) than random splits. Analysis revealed significant distribution shifts between the predefined splits: for example, 277 antibody-antigen pairs appear in training but only 6 in testing, and the test set has a lower binding rate (17.4%) than training (19.3%). This distribution imbalance, rather than model inadequacy, likely explains the performance gap, as held-out validation from the same training distribution were consistent with random-split results.

DBD5 is a well established protein-protein docking dataset used by every method we compare against. After splitting multimers into pairs as mentioned above, the resulting dataset has 335 pairs. We ran inference only, using the pre-trained PPIInterface 30% and 90% weights. The results clearly outperform BIPSPI in sequence-only mode. It achieves comparable and sometimes higher results than structure-based methods. In particular, our method achieves the highest precision out of all the tested methods. Pair-EGRET reports higher recall than precision on DBD5 (recall=0.75, precision=0.26); our method achieves more balanced precision-recall, which is generally preferable in practice.

Dockground is a dataset of 396 complexes used for protein-protein interaction and docking. As before, we split multimers into pairs, which resulted in 444 distinct pairs. We could not find any comparison to existing methods. The only method that used Dockground for benchmarking, Pair-EGRET, only tested for contact map and not interface detection.

EpiPred is an antibody-antigen dataset containing 148 proteins. We again split multimers into pairwise interactions, resulting in 286 pairs. Our method outperforms PInet in all available metrics except recall. However, the high recall obtained by PInet results in very low precision while our model is much more balanced.

As shown in our experiments on the PPInterface dataset, as well as across all five datasets evaluated, our method maintains close precision-recall balance (within ≈ ±0.1 across datasets). This balance reflects the use of MCC-optimized threshold tuning on validation, which penalizes prediction imbalance directly, in contrast to F1-optimized tuning which can produce skewed precision-recall trade-offs. The balanced behavior also means that reported metrics like F1 and MCC represent the model’s genuine ranking capability rather than artifacts of aggressive recall maximization. As seen below, it makes our model different to other methods that are tuned towards high recall at the expense of lower precision. We provide a post-processing script that allows threshold optimization for F1 or recall, if higher recall is desired at the expense of precision.

The cross-dataset results reveal three key findings. First, the five datasets yield AUROC values from 0.784 to 0.842, demonstrating stable generalization across independently curated databases – including antibody-antigen complexes (EpIPred), the canonical DBD5 docking benchmark, and the diverse Dockground dataset. Second, the difference between the 30% and 90% PPInterface models on external datasets is rather small (Dockground: +0.029 MCC; DBD5: +0.057 MCC; EpIPred: +0.028), much smaller than the internal PPInterface inflation effect of +0.145 MCC. This indicates that the rich-redundancy training advantage seen internally does not transfer cleanly to other databases – supporting the interpretation that internal inflation reflects homolog memorization within PPInterface rather than a generally more capable model. Third, our method on PPInterface 30% is similar to independently-trained MaSIF 30% AUROC (0.811 vs 0.828), confirming that the architecture and training regime are not overfit to PPInterface-specific characteristics.

#### Case study

Figure 3 shows two of the best predicted complex in the 30% dataset: (a) The crystal structure of A. aeolicus prephenate dehydrogenase, (PDB: 3GGG, Chains B and D). The predicted interface achieved MCC, precision and recall of 0.88, 0.88, and 0.98, respectively. (b) shows the unphosphorylated receiver domain of DesR in the active state (PDB: 4LE2). The predicted interface achieved MCC, precision and recall of 0.8778, 1.0, 0.794, respectively.

**Fig 3.**
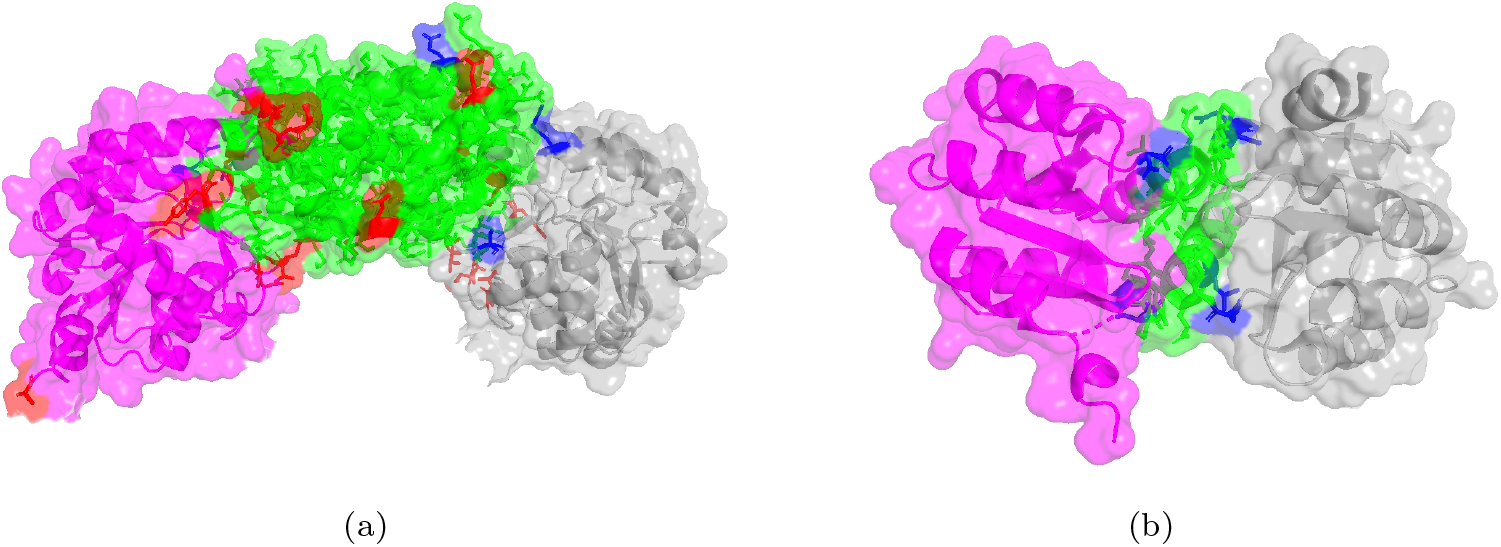
(a) The crystal structure of A. aeolicus prephenate dehydrogenase, (PDB: 3GGG, Chains B and D). (b) The crystal structure of the unphosphorylated receiver domain of DesR in the active state (PDB: 4LE2). Correctly predicted interface residues are shown in green, false positives in red and false negatives in blue.

## Discussion

Our partner-specific protein-protein interaction model was trained on a very large dataset of protein-protein pairs with known structures at different levels of redundancy. To the best of our knowledge, this is the most comprehensive training set used for this purpose. This allows us to test the effect of sequence redundancy on training large language models for protein-protein interface detection. Our primary finding is that sequence redundancy substantially inflates reported performance in partner-specific binding site prediction. Using identical model architecture and hyperparameters, the same model achieves AUROC=0.811 at 30% identity filtering versus AUROC=0.890 at 90%, a difference of +0.079 attributable entirely to evaluation rigor. The MCC and AUPRC inflation effects are larger (+0.145 MCC, +0.162 AUPRC). Beyond the absolute inflation, the per-pair efficiency analysis reveals a striking dissociation: pairs added between 50% and 90% identity contribute 4× less performance gain per pair than those added between 30% and 50% identity. This pattern can be explained by homolog memorization, rather than by genuine learning from new data.

Training and testing our model on the full and non-redundant MaSIF dataset gives near-identical evaluation metrics, hinting that the full MaSIF dataset already has rather low redundancy. Cross-dataset inference experiments on much smaller datasets confirm that the strongest redundancy inflation effect is internal to PPInterface rather than a general property of the trained model. On Dockground, switching from the 30% to 90% PPInterface model changes MCC by only +0.029. On DBD5 it changes by +0.057, and on EpIPred it increases by +0.028. This pattern strengthens our observation that much of the apparent 90%-model superiority of the PPInterface comes substantially from homolog memorization within the dataset itself – pairs the 90% training set “recognizes” but the 30% set does not – and does not generalize to externally curated benchmarks. The clean cross-dataset transfer of the 30% model is the strongest evidence that the model learns genuine, transferable sequence-to-binding-site signal. Importantly, model achieves AUROC=0.824–0.832 on independently-trained MaSIF, slightly superior to the PPInterface results, and inference AUROC values of 0.784–0.842 across the five datasets (PPInt, MaSIF, DBD5, EpIPred, Dockground). This consistency across independently curated databases, spanning general PPI, antibody-antigen, docking benchmarks, and the canonical DBD5 structural benchmark, confirms that our method learns genuine sequence-to-binding-site signal that generalizes beyond its training distribution.

To the best of our knowledge, this is the largest dataset used for this task, and the first thorough redundancy inflation study. Most published methods use small datasets with variable or no redundancy filtering, often at 90% or higher identity thresholds. Our results suggest that cross-study performance comparisons in this field should be interpreted with substantial caution. We recommend future work either report performance at multiple redundancy thresholds (allowing readers to assess the inflation effect for themselves) or adopt a consistent threshold (e.g., 30% sequence identity) to enable fair comparison across studies.

The negative-control hierarchy provides a clean decomposition of where the model’s predictive signal comes from. Globally random labels yield random AUROC=0.500, confirming no information leakage in the data pipeline. Within-chain shuffled labels yield AUROC=0.653 – a real signal floor reflecting per-chain composition statistics (chains with more interface-prone amino acids systematically contain more binding sites). The full model’s AUROC=0.811 contributed an additional genuine sequence-to-binding-site signal beyond what is learnable from chain-level statistics alone. The size-matched comparison with ESM-2 reveals a striking asymmetry. At matched ≈ 3*B* parameters with frozen encoders, ProstT5 outperforms ESM-2. Scaling ESM-2 from 650M to 3B improves MCC by only +0.019, indicating diminishing returns from model capacity alone. LoRA fine-tuning improves ProstT5 by a small but consistent amount but consistently destabilized ESM-2, regardless of hyperparameter configuration and distance convention. We hypothesize that the 3Di structural alphabet aligns the pre-training objective with the geometric properties of protein interfaces, creating representations that are both inherently more discriminative for binding site prediction and that occupy a region of parameter space where small task-specific updates further refine the signal rather than disrupt it.

Our ablation results reveal that cross-chain attention and LoRA fine-tuning are remarkably substitutable mechanisms for incorporating partner-specific information. The no-cross-attention LoRA=8 configuration (MCC=0.343) is close to the full-model LoRA=0 configuration (MCC=0.36), two architecturally distinct mechanisms (encoder fine-tuning vs cross-chain attention) arrive at a similar final performance. Removing the LoRA modification when cross-attention is present only adds +0.007 to the MCC and 0.009 to AUROC. A practically useful consequence: ProstT5 with cross-attention but no LoRa modification achieves nearly the same performance as the fully fine-tuned model at substantially lower computational cost, an attractive option for resource-constrained deployment.

The comparison across distance conventions reveals that the 6Å heavy-atom convention, matching most methods’ standard, gives the best metric profile across most measures. The 8Å *Cα* convention has slightly higher class balance (17% binding) to 6Å heavy-atom (≈ 13%) but produces noisier labels because *Cα* proximity does not always reflect side-chain interaction. This finding is itself methodologically useful: papers using 8Å *Cα* labeling are training on noisier targets than papers using heavy-atom labeling, even at rather similar class balance. 6Å *Cα* labeling is very sparse (≈ 5%) and may miss important interactions between side chains.

Several limitations should be acknowledged. Our ESM-2 LoRA comparisons did not converge cleanly across multiple distance conventions despite extensive hyperparameter sweeps; while this itself is a finding (and reproduced across three different labeling conventions), it precludes us from reporting LoRA-tuned ESM-2 numbers. The cross-dataset inference experiments use the PPInterface-trained model directly, without de-contaminating the external datasets against the PPInterface training set at 30% identity. It is almost certain that the PPInterface training weights include similar chains due to the dataset’s large size; the small difference between 30% and 90% model results across external datasets (+0.028–0.055) suggests that the model is not relying heavily on memorized homologs at inference time, but residual contamination cannot be entirely excluded. MaSIF’s predefined train/test split exhibits distribution shift (e.g., severe antibody-antigen imbalance between splits) that complicates interpretation of results on that benchmark. Finally, generalization to protein-peptide, protein-nucleic-acid, and transient signaling interactions has not been assessed.

## Conclusion

We present a sequence-based, partner-specific protein-protein binding site predictor combining ProstT5 with cross-chain attention and contact supervision, trained on a large, systematically non-redundant benchmark of ≈ 14, 000–32, 500 pairs compiled from the PPInterface database. Our analysis of redundancy inflation quantifies a methodological confound that has likely affected reported performance across the field: AUROC varies by +0.079 and MCC by +0.145 purely due to filtering stringency on the internal benchmark, with per-pair efficiency analysis isolating homolog memorization as the dominant mechanism behind the high-identity performance jumps. Cross-dataset inference on three independent benchmarks (DBD5, EpIPred, Dockground) shows that this internal inflation effect does not transfer cleanly to externally curated benchmarks: switching between the PPInterface 30% and 90% models changes external-dataset MCC by only +0.029 to +0.055, supporting the interpretation that the larger internal inflation reflects homolog memorization rather than a uniformly more capable model. At rigorous 30% sequence identity filtering, our method achieves AUROC values of 0.784–0.828 on five datasets, confirming genuine cross-dataset generalization. A size-matched comparison with ESM-2 demonstrates that structural pre-training in the encoder is an important factor for sequence-only binding site prediction, providing both substantially better representations and a more amenable fine-tuning landscape across all interface conventions tested. These results motivate further development of structurally-aware protein language models for interface prediction and related tasks, and underscore the need for rigorous redundancy filtering in benchmark evaluations. Our method outperforms existing sequence-only partner-specific binding site prediction methods and has comparable performance to many structure-based methods, despite not involving direct structural information at training and evaluation. Future work includes adding structure-based features such as solvent-accessible surface area or secondary structure elements, as well as training on more advanced large language models. Another direction involves the use of known structures as a post-processing step to improve accuracy.

## Acknowledgments

The training was performed on the UMass Boston research cluster, Chimera.

