## Supplementary material for "Handshake: Partner-Specific Protein-Protein Binding Site Prediction at Scale Using ProstT5 and Cross-Chain Attention"

Nurit Haspel[1*]

**1** University of Massachusetts Boston, Department of Computer Science, Boston MA USA

*

### Negative Controls

**Table S1** Negative-control hierarchy.

| Label condition | AUROC | MCC | Interpretation |
|---|---|---|---|
| Globally random labels | 0.500 | 0.001 | No leakage / artifacts |
| Within-chain shuffled | 0.653 | 0.161 | Per-chain composition signal |
| Real labels (full model) | 0.811 | 0.367 | Genuine binding-site signal |

**Figure S1** AUROC for Negative control experiment. 0.5 indicates random labels.

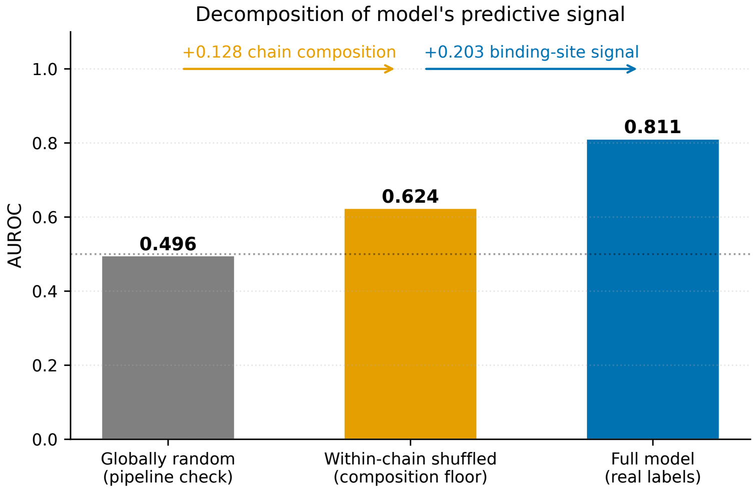


### Architectural Ablation and Comparison with ESM-2. Cross-attn = cross attention between the two interacting chains.

**Table S2** ESM-2 vs ProstT5 comparison.

| Model | Parameters | LoRA | MCC | AUROC | AUPRC |
|---|---|---|---|---|---|
| ESM-2 650M | 650M | 0 | 0.244 | 0.735 | 0.330 |
| ESM-2 3B | 3B | 0 | 0.277 | 0.754 | 0.361 |
| ProstT5, No cross-attn | 3B | 0 | 0.295 | 0.763 | 0.369 |
| ProstT5, No cross-attn | 3B | 8 | 0.343 | 0.793 | 0.431 |
| ProstT5 + Cross-attn | 3B | 0 | 0.360 | 0.802 | 0.451 |
| ProstT5 + Cross-attn | 3B | 8 | 0.367 | 0.811 | 0.462 |

**Figure S2** MCC and AUROC values for the ESM-2 650M and 3B vs ProstT5 frozen and LoRA comparison on the 30% 6Å heavy-atom dataset. Adding LoRA modification to the ESM-2 models caused their collapse.

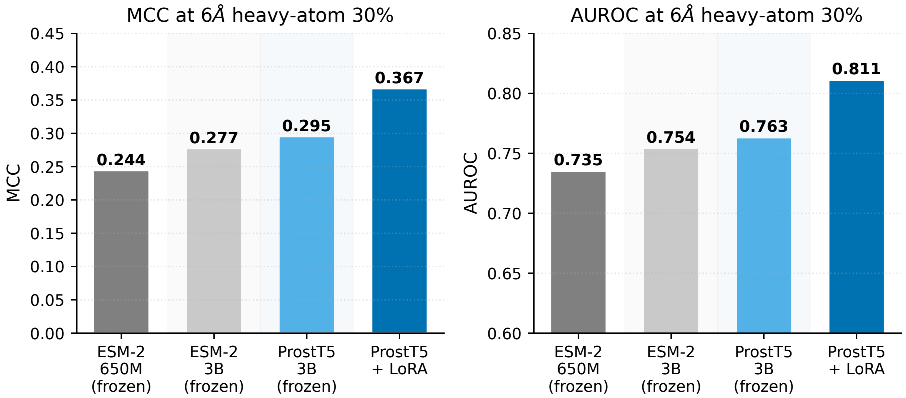


The encoder comparison holds across all three distance conventions tested. At 30% identity, ProstT5 outperforms ESM-2 3B by +0.054 MCC at 8Å $C\alpha$, +0.088 MCC at 6Å $C\alpha$, and +0.094 MCC at 6Å heavy-atom. The structural pretraining advantage is consistent and grows slightly as the labeling convention becomes cleaner.

### Comparison with Distance Conventions

We additionally trained the same model architecture on two alternative interface conventions: 6Å $C\alpha$–$C\alpha$ (a stricter labeling that includes only residues with $C\alpha$ atoms within 6Å, $\approx 5\%$ binding) and 8Å $C\alpha$–$C\alpha$ (a more permissive $C\alpha$-based labeling, $\approx 17\%$ binding). Table S4 compares the three at 30% redundancy filtering.

**Table S3** Performance under three interface conventions at 30% identity filtering. The 6Å heavy-atom gives the best metric profile across most measures.

| Convention | Binding % | MCC | AUROC | AUPRC | F1 |
|---|---|---|---|---|---|
| 6Å heavy-atom (PPInt) | 13% | 0.373 | 0.811 | 0.462 | 0.485 |
| 6Å $C\alpha$ (strict) | 5% | 0.316 | 0.827 | 0.320 | 0.347 |
| 8Å $C\alpha$ (permissive) | 17% | 0.264 | 0.714 | 0.383 | 0.347 |

The 6Å heavy-atom convention dominates on MCC, AUPRC, and F1, while the 6Å $C\alpha$ convention has higher AUROC despite lower MCC (a quirk of the very sparse positive class producing well-ranked but conservative predictions). The 8Å $C\alpha$ convention, similar in class balance to 6Å heavy-atom, underperforms across all metrics, indicating that 8Å $C\alpha$ labels are noisier than 6Å heavy-atom labels even at comparable density. This is consistent with the heavy-atom convention capturing genuine side-chain interface contacts while the 8Å $C\alpha$ convention conflates real interfaces with mere proximity.

**Table S4 Performance across redundancy thresholds (6Å $C\alpha$–$C\alpha$ dataset).** Stricter $C\alpha$-based labeling at 6Å, $\approx 5\%$ binding rate. Lower MCC and AUPRC than the heavy-atom convention but the highest AUROC, reflecting the very sparse positive class producing well-ranked predictions.

| Dataset | Pairs | MCC | F1 | AUROC | AUPRC |
|---|---|---|---|---|---|
| 30% (MMseqs2) | 11,327 | 0.316 | 0.347 | 0.827 | 0.320 |
| 50% (CD-HIT) | 13,606 | 0.368 | 0.396 | 0.855 | 0.371 |
| 90% (CD-HIT) | 27,071 | 0.439 | 0.463 | 0.895 | 0.456 |

**Table S5 6Å $C\alpha$ ablation results on the 30% dataset.** Three-way ablation including the no-supervision configuration, complementing the 2×2 ablation in the main paper. Format: MCC / AUROC.

| Configuration | LoRA=0 (frozen) | LoRA=8 (tuned) |
|---|---|---|
| No cross-attention | 0.231 / 0.781 | 0.274 / 0.805 |
| Cross-attn, no contact supervision | 0.294 / 0.809 | 0.305 / 0.822 |
| Full model | 0.305 / 0.818 | 0.316 / 0.834 |

**Table S6 Performance across redundancy thresholds (8Å $C\alpha$–$C\alpha$ dataset).** Permissive $C\alpha$-based labeling at 8Å, ≈ 17% binding rate. Underperforms 6Å heavy-atom across all metrics despite similar class balance, indicating 8Å $C\alpha$ labels are noisier than heavy-atom labels.

| Dataset | Pairs | MCC | MCC (+prop) | AUROC | AUPRC |
|---|---|---|---|---|---|
| 30% (MMseqs2) | 14,220 | 0.264 | 0.269 | 0.714 | 0.383 |
| 50% (CD-HIT) | 18,312 | 0.289 | 0.299 | 0.737 | 0.409 |
| 90% (CD-HIT) | 32,992 | 0.345 | 0.347 | 0.779 | 0.463 |

**Table S7 8Å ablation results on the 50% dataset.** Pattern matches the 6Å ablations: cross-chain attention and LoRA are partially substitutable, with the largest LoRA contribution observed when cross-attention is absent. Format: MCC after confidence propagation.

| Configuration | LoRA=0 | LoRA=8 |
|---|---|---|
| No cross-attention | 0.234 | 0.277 |
| Cross-attn, no contact supervision | 0.277 | 0.290 |
| Full model | 0.287 | 0.299 |

**Table S8 8Å ESM-2 vs ProstT5 comparison on the 50% dataset.** Unlike at 6Å, ESM-2 LoRA training was stable at 8Å but consistently underperformed the frozen baseline due to LoRA-induced collapse toward the majority class.

| Model | Parameters | LoRA | MCC | AUROC | AUPRC |
|---|---|---|---|---|---|
| ESM-2 650M, frozen | 650M | 0 | 0.222 | 0.679 | 0.346 |
| ESM-2 3B, frozen | 3B | 0 | 0.235 | 0.689 | 0.356 |
| ESM-2 650M + LoRA | 650M | 8 | 0.171 | 0.628 | 0.299 |
| ESM-2 3B + LoRA | 3B | 8 | 0.174 | 0.626 | 0.300 |
| ProstT5, frozen | 3B | 0 | 0.276 | 0.724 | 0.397 |
| ProstT5 + LoRA | 3B | 8 | 0.289 | 0.737 | 0.409 |

### Contact Map Prediction

As a secondary output, our model predicts residue-residue contact maps across the interface. Notably, in the 6Å heavy-atom configuration the contact head's predictions become sufficiently discriminative that combining contact-map evidence with sequence-neighbor evidence (`require_both=True`) is the optimal post-processing configuration, unlike the 6Å $C\alpha$ and 8Å $C\alpha$ runs where sequence-neighbor evidence alone was sufficient. This indicates the 6Å heavy-atom labeling produces clean enough contact supervision for the contact head to provide independent confirmation of binding-site predictions.

**Table S9 Contact map prediction AUPRC within predicted binding sub-matrix** (bind_thresh=0.3, 8Å datasets where contact pairs were generated).

| Dataset | Contact AUPRC | Random baseline | Lift over random |
|---|---|---|---|
| 50% (8Å) full model | 0.072 | 0.0017 | $\approx 43\times$ |
| 70% (8Å) full model | 0.068 | 0.0016 | $\approx 44\times$ |
| 90% (8Å) full model | 0.155 | 0.0015 | $\approx 101\times$ |

**S1 File. Source code, data and pre-trained model weights.** Available at `http://www.github.com/nurith/Handshake`.
